# Measuring bridging forces in protein-DNA condensates

**DOI:** 10.1101/2025.01.29.635494

**Authors:** Vikhyaat Ahlawat, Hashini Ekanayake Mudiyanselage, Huan-Xiang Zhou

## Abstract

Protein-DNA condensates mediate transcription and regulate gene expression and DNA replication and repair. The intermolecular bridging forces stabilizing condensates have direct roles in these processes. Here we use optical tweezers to measure bridging forces. In the presence of protamine, a single condensate is observed on a 20.5-knt single-stranded DNA (ssDNA) tethered between two microbeads. Stretching produces force curves with a sawtooth pattern, suggesting that the condensate is dissembled by the sequential rupture of individual protamine-ssDNA bridges. The bridging forces are 11.3 ± 4.6 pN, with unfolding lengths of 1.3 ± 0.8 µm for single bridges. In contrast, double-stranded DNA (dsDNA) forms protamine-bridged tangles that can withstand forces high enough (∼55 pN) for strand separation. ssDNA tracks unpeeled at nicks on dsDNA by overstretching seed tangle formation upon retraction, but the initial condensates have a sufficient ssDNA-to-dsDNA ratio to appear liquid-like, as indicated by a sawtooth pattern in the subsequent stretching. The presence of dsDNA raises bridging forces to 34 ± 8 pN, which revert to ∼10 pN upon adding external ssDNA. In line with these single-molecule results, protamine-dsDNA mixtures form solid-like aggregates and require the addition of ssDNA to become liquid droplets. Conversely, adding dsDNA slows the fusion of protamine-ssDNA droplets. This work demonstrates the first measurements of bridging forces and shows that the ssDNA-to-dsDNA ratio can tune their magnitude in protein-DNA condensates.

## Introduction

Protein-DNA condensates mediate transcription ^1, 2^ and regulate gene expression ^3–5^ and DNA replication ^6^ and repair ^7–12^. Condensation helps the recruitment of factors and facilitates the assembly of functioning complexes. Such factors include molecular motors that drive these DNA-related processes, e.g., RNA polymerases for transcription, replicative helicases and DNA polymerases for replication, and repair helicases for repair. Single-molecule studies have shown that DNA-dependent motors can work against an impeding force ^13^. For example, *Escherichia coli* RNA polymerase stalled only at forces of ∼15-25 pN ^14, 15^; *Saccharomyces cerevisiae* RNA polymerase II ceased to transcribe at 7.5 pN force ^16^. In comparison, the stall forces of Klenow and T7 DNA polymerases were 20-35 pN ^17, 18^. For RecG and UvsW, a 35 pN force only moderately slowed the DNA rewinding rate, but a further increase in force resulted in the dissociation of the DNA repair helicases from the single-stranded (ss) and double-stranded (ds) junction ^19^. Impeding forces may be presented by other DNA-bound proteins and by higher-order structures of the DNA molecule.

Protamine is a small, Arg-rich intrinsically disordered protein (IDP) that, in the sperm nucleus, replaces histones to further compact chromatin by >10-fold. Whereas histones compact somatic chromatin by forming nucleosomes, the mechanism of sperm chromatin compaction by protamine is still unresolved ^20, 21^. During the histone-to-protamine transition, numerous DNA strand breaks occur in the spermatid nucleus to relieve supercoils as histones are removed ^22, 23^. Protamine expression also starts at this stage. As spermatids mature into sperm, DNA strand breaks are repaired and transcription is globally silenced.

Our recent optical tweezers (OT) study uncovered a piece of the sperm chromatin puzzle ^24^. Salmon protamine, with the sequence M_1_PRRRRSSSR P_10_VRRRRRPRV S_20_RRRRRRGGR R_30_RR, condenses a single dsDNA chain into “tangles” that withstand forces strong enough (∼55 pN) to separate the two strands of DNA, along with bends and loops that unravel at 10-40 pN forces. We assumed that tangles were bundles of loops bridged by protamine molecules. As their strength far exceeds the stall force (7.5-25 pN) of RNA polymerases ^14–16^, protamine-bridged tangles provide a mechanism for global transcription silencing. We also found that, when dsDNA was overstretched to produce strand separation at nick sites, tangles emerged at ssDNA-dsDNA junctions. One consequence of strand separation is the exposure of nucleobases, allowing for easy access by Arg sidechains. In line with that, our molecular dynamics simulations further revealed that Arg sidechains not only form salt bridges with the phosphate backbone of DNA but also frequently hydrogen bonds with nucleobases from both strands simultaneously. These Arg wedges formed with dsDNA explain the hyper-stability of tangles.

In addition to protamine, DNA condensation by several other proteins has been studied by OT or magnetic tweezers. Condensates were induced by pioneer transcription factors FoxA1 ^25^ and Klf4 ^26^ and the nuclear enzyme PARP1 ^27^ on single λ-DNA (48.5 kb) molecules under a low force. By contrast, condensates induced by pioneer transcription factor Sox2 and heterochromatin-associated protein HP1α on λ-DNA were stable at least to 40 pN ^28^ and to strand separation ^29^, respectively. Similarly, an 8.1-kb DNA condensed by the HIV-1 nucleocapsid protein was stable at least to 50 pN ^30^. In comparison, linker histone H1 ^31^ and prion-like protein FUS ^32^ formed co-condensates only with unpeeled ssDNA tracks at nick sites of overstretched λ-DNA. Condensates induced by poly-L-lysine on single ssDNA molecules were also stable up to 50 pN force ^33^. In some cases, protein-induced DNA condensation was indicated by an increase in the force baseline when a DNA tether at low extension was moved into the protein channel. Baseline shifts of 0.5 to 9.5 pN have been observed ^25, 29, 30, 33^.

Here we used OT and brightfield and confocal microscopy to characterize condensates formed by binary mixtures of salmon protamine with ssDNA and ternary mixtures of protamine with ssDNA and dsDNA. Whereas the mixtures of protamine with dsDNA form aggregates ^24^, its mixtures with ssDNA form liquid droplets, similar to previous observations with poly-L-lysine as the protein component ^34, 35^. In the presence of protamine, a single condensate is observed on a 20.5-knt single-stranded DNA tethered between two optically trapped microbeads. Stretching produces force curves with a sawtooth pattern, suggesting that the condensate is dissembled by the sequential rupture of individual protamine-ssDNA bridges. The bridging forces are 11.3 ± 4.6 pN, with unfolding lengths of 1.3 ± 0.8 µm for single bridges. Likewise, condensates initiated by ssDNA tracks unpeeled at nicks on λ-DNA appear liquid-like, as indicated by a sawtooth pattern in the subsequent stretching. The presence of dsDNA raises bridging forces to 34 ± 8 pN, which revert to ∼10 pN upon adding external ssDNA. This work demonstrates the first measurements of bridging forces and shows that the ssDNA-to-dsDNA ratio can tune their magnitude in protein-DNA condensates.

## Results

### Protamine drives condensate formation on partially unpeeled DNA

We tethered a 17.853-kb dsDNA between two microbeads (4.35 µm diameter) to study protamine-DNA condensates in a single-molecule setting. This dsDNA construct has two defined nicks 5.005-knt apart on one strand; this 5.005-knt track of ssDNA can be peeled off by overstretching (to ∼60 pN). After staining with the intercalation dye Sytox Orange, a confocal image of the DNA molecule before the peeling off of the 5.005-knt track shows continuous staining between the two beads, but the counterpart after the peeling off shows a gap indicating that only one strand is left between the two defined nicks (Figure 1A). We refer to the two versions of the DNA molecule “intact” and “gap”, respectively. In the buffer channel (10 mM pH 7 imidazole with 150 mM KCl), the force-extension curve of the intact DNA molecule displays a slow initial rise, reaching 1 pN only at an extension of 5.2 µm, followed by a sharp rise, to a contour length of ∼6.1 µm at high force (Figure 1B). This behavior is expected of dsDNA with a persistence length of 50 nm. In comparison, the entire force-extension curve of the gap DNA molecule displays only a gradual rise. Both the faster initial rise and the longer final contour length can be attributed to the ssDNA track of the gap DNA molecule, which has a much shorter persistence length (∼ 1 nm) and has approximately twice the contour length per nucleotide compared to dsDNA (0.66 nm per nucleotide vs. 0.34 nm per base pair).

**Figure 1.**
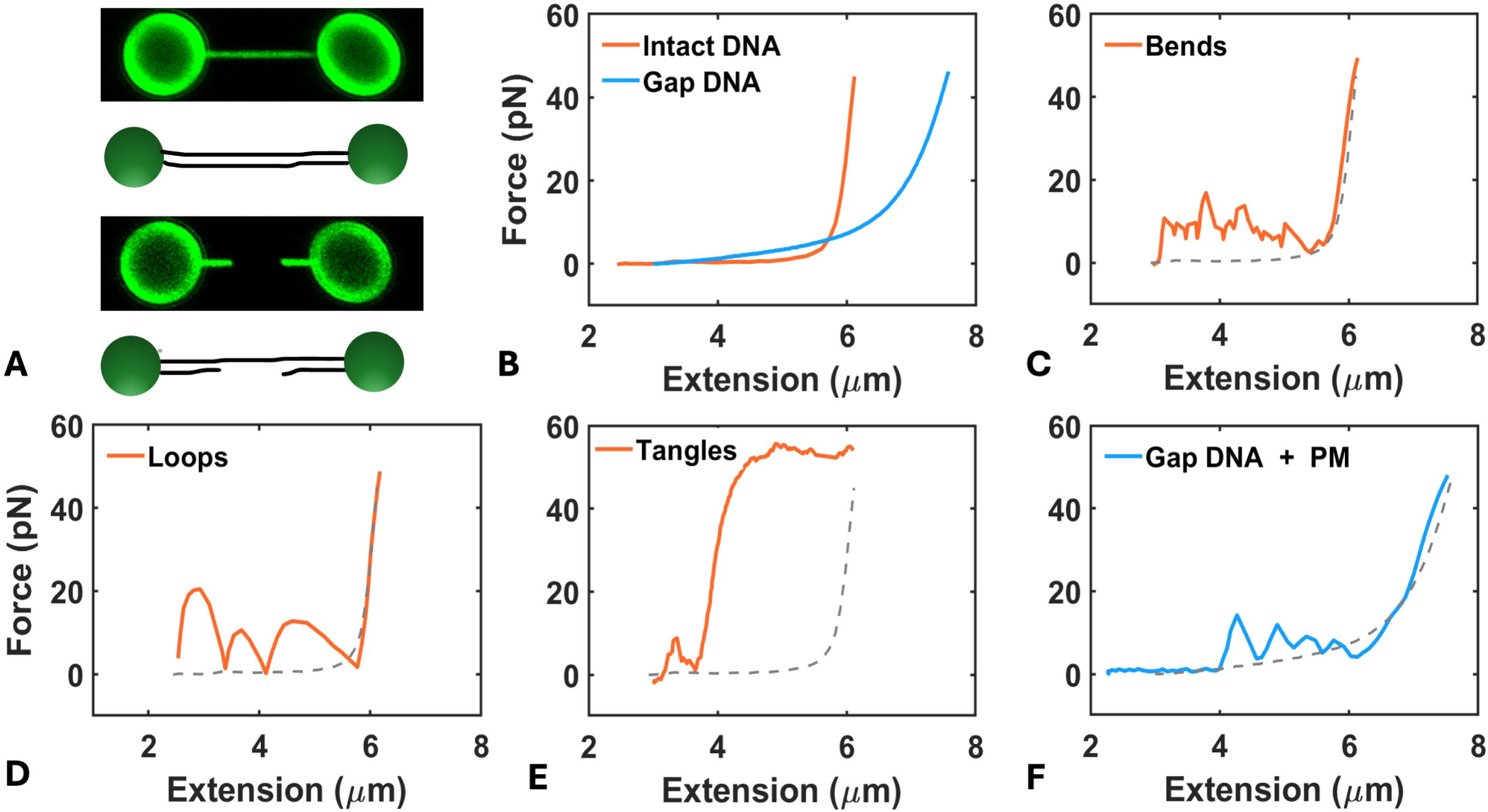
2D scans and force-extension curves of intact and gap DNA. (A) Top: 2D scan of and schematic illustrating intact dsDNA (17.853 kb); bottom: “gap DNA” configuration in which the middle 5.005-knt ssDNA track on one strand is peeled off. Sytox Orange (0.5 μM) was used for staining dsDNA. (B) Force-extension curves of intact and gap DNA in the buffer channel. (C-E) Force-extension curves of intact DNA in the protamine channel, showing bending, looping, and tangling modes, respectively. (F) Force-extension curve of gap DNA in the protamine channel, showing a sawtooth pattern implicating sequential unraveling of a liquid condensate. In (C-F), the baseline was shifted after transferring the retracted tether from the buffer channel to the protamine channel (Figure S1); the baseline was reset to 0 before stretching. Also, the respective stretching curve in buffer was displayed as a dashed curve for reference.

We have shown that protamine compacts dsDNA in three modes, with distinct patterns in force-extension curves ^24^. When the retracted intact DNA tether (extension at 2-3 µm) is transferred from the buffer channel to the protamine channel (10 µM), a sudden rise of ∼10 pN in the baseline force is observed (Figure S1A). Such a baseline shift has been observed previously ^25, 29, 30, 33^. We reset the baseline to 0. The subsequent stretching curves of the intact DNA molecule exhibit the three known modes (Figure 1C-E). The bending mode is characterized by rapid small-amplitude fluctuations around 10 pN of pulling force (Figure 1C) and arises from protamine-induced local bends. In the looping mode, the pulling force rises quickly to ∼20 pN but then falls slowly, with a smooth turnover at the force peak (Figure 1D); the fall in force corresponds to the rupture of a protamine-bridged loop. The tangling mode is indicated by an early rise to the force plateau at ∼55 pN corresponding to strand separation (Figure 1E), meaning that the protamine-mediated structure is stable enough to withstand such a high force.

The gap DNA also displays a baseline shift upon transferring to the protamine channel. However, the subsequent stretching curves consistently exhibit a sawtooth pattern (Figure 1F), indicating a new condensation mode. The peak forces of the new mode are ∼10 pN, similar to those in the bending mode Figure 1C, but the increase in extension in a single rupture event is much greater (∼0.1 µm in Figure 1C vs. 0.5-1 µm in Figure 1F). The bridging forces in this new mode are softer than those in the looping mode of Figure 1D, as indicated both by the lower peak forces and by a slower rise and a faster fall in each rupture event. The new mode is associated with ssDNA because only the gap DNA, with a 5.005-knt ssDNA track in the middle of the molecule, produces this mode. The dsDNA segments flanking the ssDNA track likely bind protamine and thus can form bends. The small increases in extension by the rupture of these bends might be superimposed on the much larger increases in extension from the rupture of protamine-ssDNA bridges. To avoid such complications due to the simultaneous presence of ssDNA and dsDNA on the same molecule, we next turn to a pure ssDNA tether.

### Protamine-ssDNA condensates exhibit sequential unraveling under force

We produced a 20.452-knt ssDNA tether between two microbeads (Figure 2A). After staining with SYBR Gold, continuous 2D scans were performed (1.45 s per frame) while the ssDNA was pulled at 1 µm/s in the protamine channel from an initial extension of 3 µm (Figure 2B). A condensate is formed on the ssDNA molecule; its size decreases progressively, suggestive of sequential unraveling.

**Figure 2.**
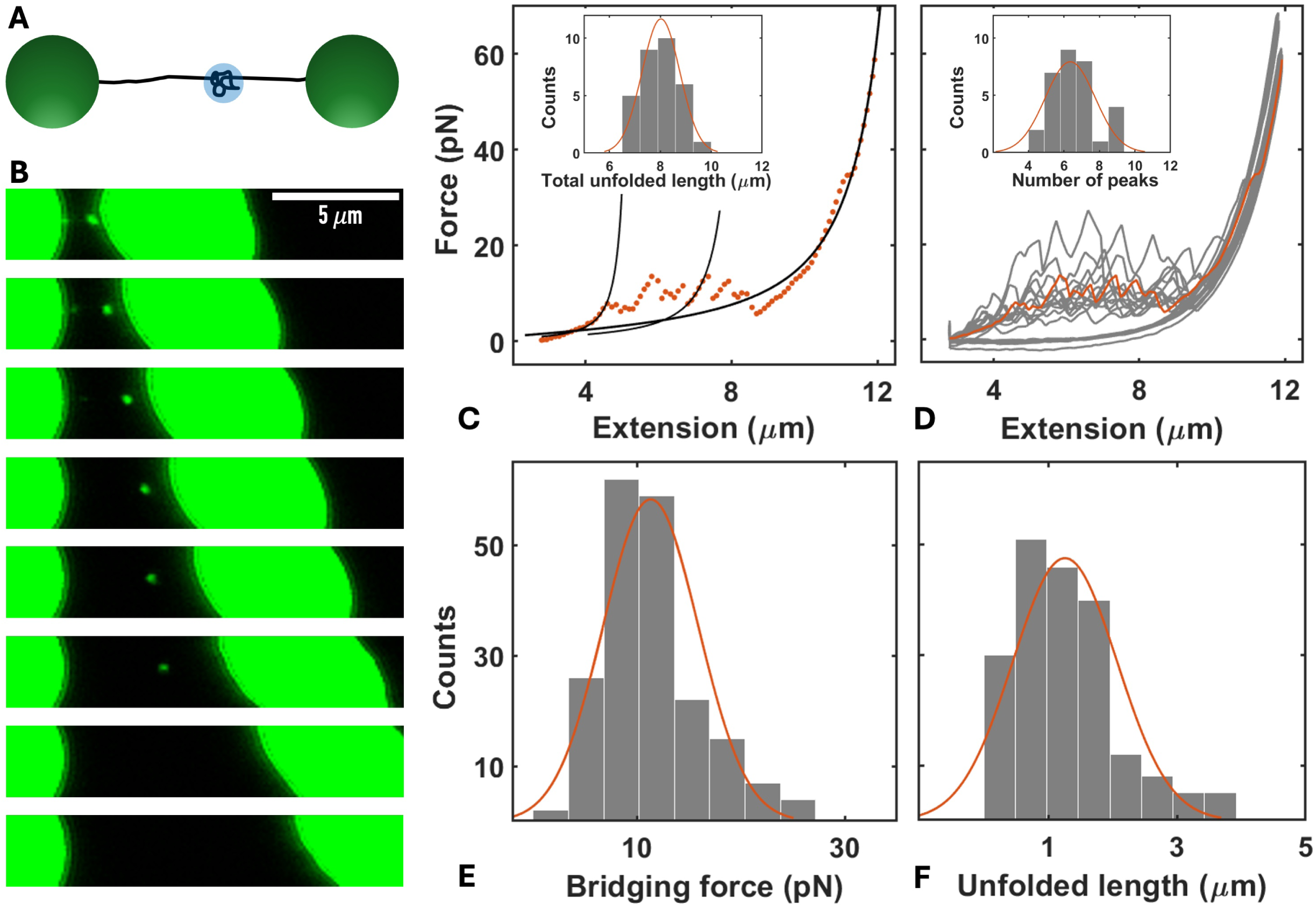
2D scans and force-extension curves showing unraveling of protamine-ssDNA condensates. (A) Schematic illustrating a ssDNA tether with a condensate formed in the middle. (B) Continuous 2D scans of a 20.452-knt ssDNA tether during stretching in the protamine channel containing 0.5 µM SYBR Gold. The left bead was fixed while the right bead was pulled at 1 µm/s from an initial extension of 3 µm. The right bead appears distorted because each frame takes a finite time (1.45 s). The condensate is unraveled by the 7th frame, where the extension reaches ∼10 µm. (C) Force-extension curve showing a sawtooth pattern of rupture of individual protamine-ssDNA bridges. Fits of the first, a middle, and the final rises in force to the wormlike-chain model are displayed as black curves. Inset: histogram of the total increased contour length after a condensate is completely unraveled. (D) Reversible condensate formation demonstrated by reproducible sawtooth patterns in 14 successive stretch-retract cycles; the first of these stretching curves, already shown in (C), is displayed in orange color. Inset: histogram of the number of force peaks per stretching curve. (E) Histogram of individual bridging forces. (F) Histogram of the increased contour length upon the rupture of a single bridge. In (C-F), curves superimposed on bar graphs display Gaussian fits.

The ssDNA again displays a baseline shift upon transferring to the protamine channel (Figure S1B). The subsequent stretching curve exhibits the sawtooth pattern already seen in Figure 1F (Figures 2C and S2). The first and last force peaks typically occur at 4-5 µm and 8-9 µm of extensions, respectively. After stretching to 12 µm, the subsequent retraction (to 3 µm) conforms to a wormlike chain model ^36^, but the next stretching curve again exhibits the sawtooth pattern (Figures 2D). Indeed, the foregoing behavior repeats itself over many stretch-retract cycles, suggesting that after the rupture of protamine-ssDNA bridges, protamine molecules are still bound to the ssDNA and ready to reform bridges upon the retraction of the ssDNA. Fitting of the final rise in force to the wormlike chain model yielded a contour length of 13.53 ± 0.08 µm (mean ± SD; *N* = 31), which corresponds to the expected 0.66 nm per nucleotide. The resulting persistence is 1.2 ± 0.1 nm. Fitting the first rise in force to the wormlike chain model yielded a contour length of 5.5 ± 1.6 µm, which mostly comes from the sequences flanking the condensate. The difference in contour length between the final and first rises in force is the total unfolding length of the condensate. The total unfolding lengths are 8.0 ± 0.7 µm (Figure 2C inset); the mean value corresponds to 59% of the total contour length. The persistence length from fitting the first rise in force is 5.6 ± 1.6 nm, significantly longer than the counterpart for the final rise in force. The longer apparent persistence length could be caused by an expansion of the condensate under force.

The number of force peaks per stretch is 6.4 ± 1.4 (Figure 2D inset). These force peaks correspond to the rupture of protamine-ssDNA bridges and will be called bridging forces. Among 197 rupture events, the bridging forces are 11.3 ± 4.6 pN (Figure 2E). By fitting the rises to these force peaks to the wormlike chain model (Figures 2C and S2), we obtained the unfolded lengths of single bridges as the differences in contour length between successive rupture events. The unfolded lengths of single bridges are 1.3 ± 0.8 µm (Figure 2F). The persistence lengths from fitting the middle rises in force (i.e., between the first rise and the final rise) are 2.9 ± 1.5 nm, which are intermediate between those from fitting the first and final rises in force.

Whereas condensates formed on ssDNA show a sawtooth pattern in force-extension curves with peak forces of ∼10 pN, tangles formed on dsDNA withstand forces (∼ 55 pN) that are strong enough to separate the two strands of dsDNA. Based on these differences, we describe tangles as solid-like and protamine-ssDNA condensates as liquid-like.

### Bridging forces can be tuned by ssDNA-to-dsDNA ratio

ssDNA tracks unpeeled from nicks on dsDNA by overstretching seed protamine-mediated tangle formation upon retraction ^24^. We reasoned that the initial condensates may have a high ssDNA content to be liquid-like. To test this idea, we overstretched λ-DNA to ∼24 µm in buffer and then retracted it in the protamine channel either to 6-7 µm to allow for tangle formation (Figure S3) or only to 12-13 µm before tangles were fully formed (Figure 3). Overstretched λ-DNA shows one or more gaps in 2D scans (top row in Figures 3A and S3A), indicating ssDNA unpeeling from nicks and free ends (Figures 3B and S3B). Upon retraction, condensates emerge at the edges of the gaps and thus likely involve three components: protamine, ssDNA, and dsDNA. When the retraction proceeds down to 6-7 µm in extension, the condensate grows in size, suggesting that more and more dsDNA is drawn in to form a tangle (Figure S3A). In contrast, the size of the initial condensate remains nearly constant when the retract stops at 12-13 µm in extension (Figure 3A).

**Figure 3.**
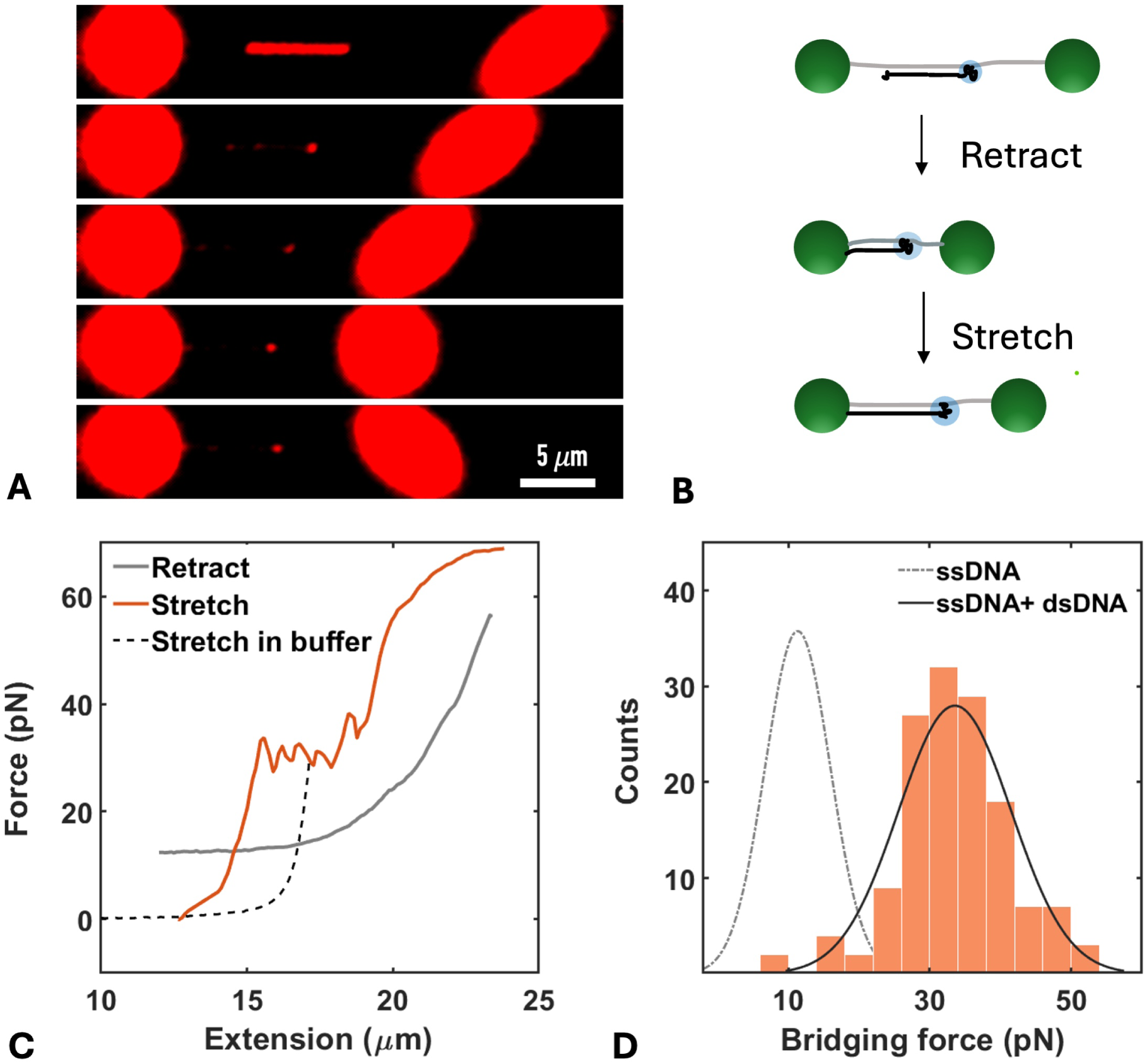
2D scans and force-extension curves of λ-DNA containing protamine-bridged condensates initiated by overstretching. (A) Continuous 2D scans during the retraction of overstretched λ-DNA to 12.5 µm and subsequent stretching. Initially, two gaps representing ssDNA tracks are observed; subsequently, a condensate emerges at the edge of the major gap. The retract-stretch cycle was performed in the protamine channel containing 1.2 µM Yoyo-3. The scan time per frame was 3.5 s. (B) Corresponding schematic illustrating the formation of a protamine-dsDNA-ssDNA ternary condensate. (C) Force-extension curves for the protocol in (A) but in the absence of the dye. The initial retraction curve (solid gray) has a baseline shift from the stretching curve in buffer (dashed gray). The baseline is reset to 0; the next stretch exhibits a sawtooth pattern (orange curve). Subsequent stretching curves are shown in Figure S4A. (D) Histogram of bridging forces from stretching curves exemplified by (C); a Gaussian fit is displayed as a solid curve. The Gaussian fit from Figure 2E is displayed as a dashed curve to demonstrate the increase in bridging force by the presence of dsDNA.

To quantify the contrasting behaviors resulting from different extents of retraction, we acquired force-extension curves (in the absence of dye). The retraction curve lags behind the stretching curve in buffer (Figures 3C and S3C), confirming ssDNA unpeeling ^37^. When the retraction proceeds to below 12 µm in extension, a sudden drop in force occurs (indicated by an arrow in Figure S3C), which is a signature of tangle formation ^24^. The force then stays at a baseline as the retraction proceeds further down to 6-7 µm in extension. This baseline is shifted by ∼10 pN from that in the stretching curve in buffer, similar to that noted for the other DNA constructs. The subsequent stretching curve exhibits the expected tangling mode, with a rapid rise to a ∼55 pN plateau much earlier than the stretching curve in buffer.

In contrast to the stretching curve after the retraction proceeds to 6-7 µm in extension, the stretching curve after a retraction terminated at 12-13 µm shows a sawtooth pattern (Figures 3C and S4A). The sawtooth pattern confirms that the initial condensates seeded by unpeeled ssDNA tracks are indeed liquid-like. Among 140 rupture events, the bridging forces are 34 ± 8 pN (Figures 3D and S4B). The latter forces are three-fold higher than the counterparts for condensates formed between pure ssDNA and protamine, demonstrating that the presence of dsDNA raises bridging forces.

If indeed it is the relatively high ssDNA-to-dsDNA ratio in the initial condensates that is responsible for the sawtooth pattern, i.e., liquid-like behavior, then the bridging forces may be reduced by the addition of external ssDNA. We thus followed a variant of the protocol in Figure 3A, where the λ-DNA tether, after retraction to 12-13 µm in the protamine channel, was soaked in a 32-nt ssDNA channel before returning to the protamine channel to perform stretch-retraction cycles (Figure S5A). In continuous 2D scans of the subsequent stretch-retract cycle, colocalized dsDNA foci (stained with YOYO-3) and ssDNA foci (labeled with Cy3) appear at the edges of unpeeled ssDNA tracks on λ-DNA tethers (Figures S5B and S6A), confirming partitioning of the external ssDNA in the condensates. For some tethers, the first stretching curve after returning to the protamine channel from the 32-nt ssDNA channel (top panel in Figure S5C) shows bridging forces at ∼35 pN, similar to those in the absence of external ssDNA (Figure 3C, D). However, subsequent stretching curves (Figure S5C) show a reduction of bridging forces to ∼10 pN, similar to those obtained on the 20.452-knt ssDNA tethers (Figure 2E). The transition from the first to the subsequent stretching curves suggests a rearrangement of ssDNA and dsDNA inside the condensates. For other λ-DNA tethers soaked with 32-nt ssDNA, the first stretching curve already shows a reduction of bridging forces to ∼10 pN (Figure S6B). Together, the results from λ-DNA tethers without and with external ssDNA demonstrate that the ssDNA-to-dsDNA ratio can tune the magnitude of bridging forces in protamine-DNA condensates.

### A sufficient ssDNA-to-dsDNA ratio is required to maintain liquidity of protamine-DNA condensates

We further carried out bulk experiments to probe the ssDNA-to-dsDNA ratio on the material properties of protamine-DNA condensates. Binary mixtures of protamine with a 32-nt ssDNA always form liquid droplets (Figure 4A), which readily fuse on contact. In contrast, binary mixtures of protamine with a 25-bp dsDNA only form amorphous aggregates; this observation remains true even when a small amount of ssDNA is added (Figure 4B). These condensates have irregular shapes and do not fuse upon coming into contact (orange arrow in Figure 4B). However, after adding more ssDNA, the condensates become liquid droplets (Figure 4C). The partition coefficient of dsDNA is 510; those of ssDNA are even higher, reaching 850-1230 (Figure 4A, C).

**Figure 4.**
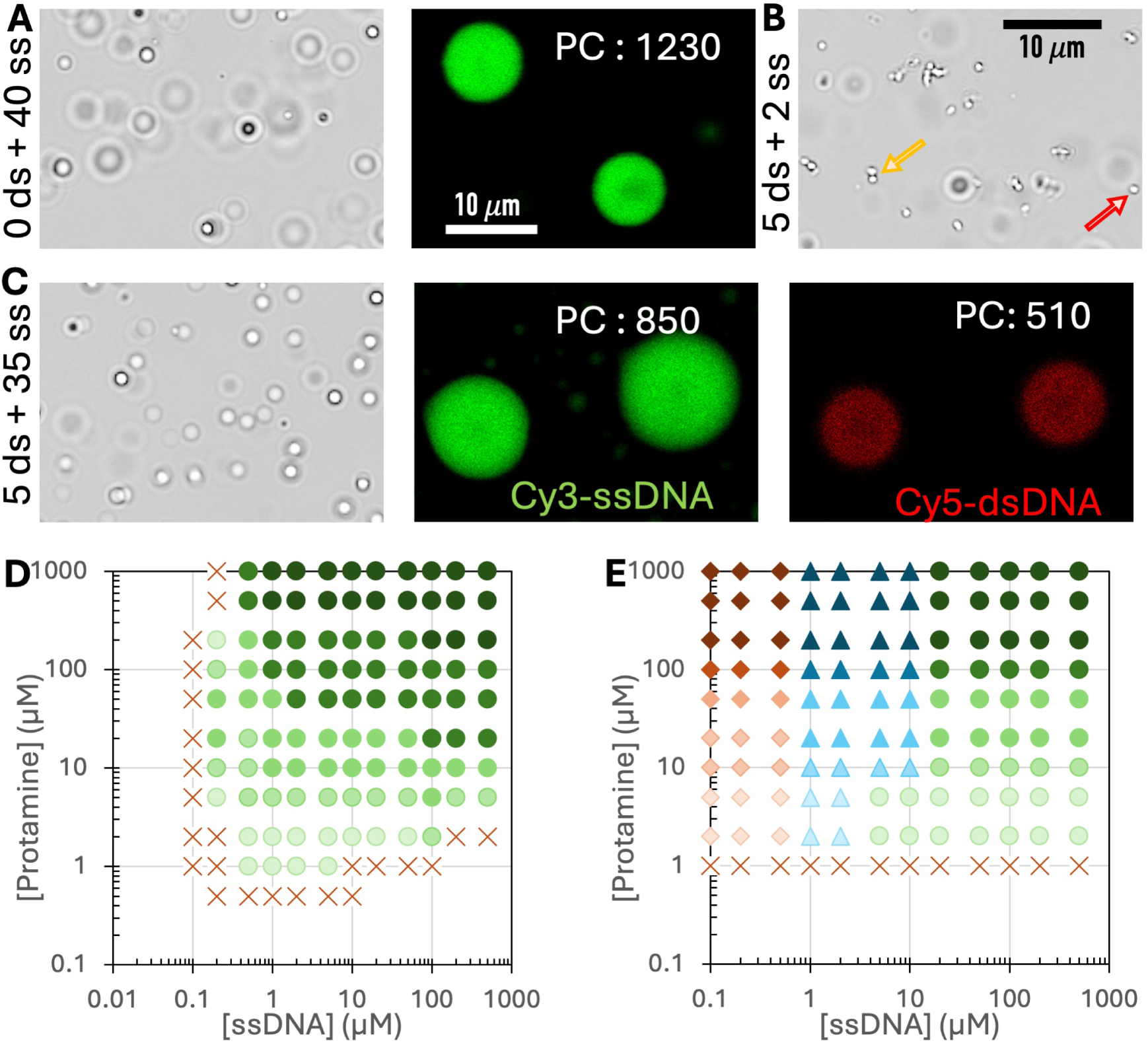
Brightfield and confocal images of protamine-DNA condensates and phase diagrams of protamine-DNA mixtures. (A) Droplets formed by mixing 40 µM protamine with 40 µM 32-nt ssDNA. (B) Aggregates formed by mixing 40 µM protamine with 5 µM 25-bp dsDNA and 2 µM 32-nt ssDNA. Aggregates can associate with each other but do not fuse (orange arrow). A small number of droplets (red arrow) also appear. (C) Droplets formed by mixing 40 µM protamine with 5 µM 25-bp dsDNA and 35 µM 32-nt ssDNA. For confocal imaging shown in (A) and (C), 0.01 µM of ssDNA or dsDNA was replaced by a variant with covalently linked Cy3 or Cy5. (D) Phase diagram of protamine and 32-nt ssDNA. (E) Phase diagram in the presence of 5 µM 25-bp dsDNA. For (D-E), crosses, circles, diamonds, and triangles indicate absence of phase separation, droplets, aggregates, and mixtures of aggregates and droplets, respectively; color intensity indicates increase in condensate number and/or size.

We determined the phase diagrams of protamine-ssDNA mixtures in the absence or presence of 2, 5, or 10 µM dsDNA. Protamine-ssDNA mixtures start to form droplets at 1 µM protamine and 0.2 µM ssDNA; droplets increase in number and/or size upon increasing the component concentrations (Figure 4D). In the presence of 5 µM dsDNA, condensates appear to be amorphous aggregates up to 0.5 µM ssDNA. With a further increase in ssDNA, some droplets appear (red arrow in Figure 4B) among aggregates. Starting at 5 µM ssDNA, condensates appear to be pure droplets at protamine concentrations up to 5 µM and again a mixture of droplets and aggregates at higher protamine concentrations. At 20 µM ssDNA, condensates are pure droplets even at the highest protamine concentration measured (1000 µM). The boundary between aggregate and droplet regions was ascertained by both observations on a brightfield microscope (irregular vs spherical shapes) and attempts to fuse two condensates on an OT instrument.

The phase diagrams in the presence of 2 and 10 µM dsDNA are similar to that at 5 µM dsDNA (Figure S7). A main difference is that the ssDNA concentration at which condensates start to be pure droplets is changed from 5 µM to 2 and 10 µM, respectively. So evidently, a minimum of a 1:1 molar ratio between ssDNA and dsDNA is required to maintain the liquidity of protamine-DNA condensates. Amorphous aggregates generally have stronger intermolecular interactions than liquid droplets ^38^. The ability of Arg sidechains to form wedges with dsDNA (as revealed by molecular dynamics simulations ^24^) but not with ssDNA may explain the stronger interaction of protamine with dsDNA than with ssDNA.

### Fusion speed of protamine-ssDNA droplets is slowed by adding dsDNA

Because protamine-ssDNA droplets readily fuse whereas protamine-dsDNA aggregates do not fuse, we can expect that doping of dsDNA into protamine-ssDNA droplets slows down fusion. Fusion speed can be accurately measured by an OT-directed method, where two droplets are trapped with low laser power and brought into contact to allow for spontaneous fusion (Figure 5A) ^39, 40^. Fusion progress can be monitored by the small residual force, which upon fitting to a stretched exponential function of time (Eq [1]) yields the fusion time 𝜏_fu_ (Figure 5B). The fusion time is proportional to the initial droplet radius 𝑅 (Figure 5C, D); the ratio, 𝜏_fu_/𝑅, is the inverse fusion speed. For droplets formed with 100 μM protamine and 70 μM 32-nt ssDNA, the inverse fusion speed is 0.90 s/μm. When 1/4 of the ssDNA is replaced with 25-bp dsDNA, the inverse fusion speed is increased to 1.14 s/μm. This increase corresponds to a 27% slowdown in fusion and can be attributed to a strengthening of protamine-DNA interactions by the replacement of a portion of the ssDNA with dsDNA.

**Figure 5.**
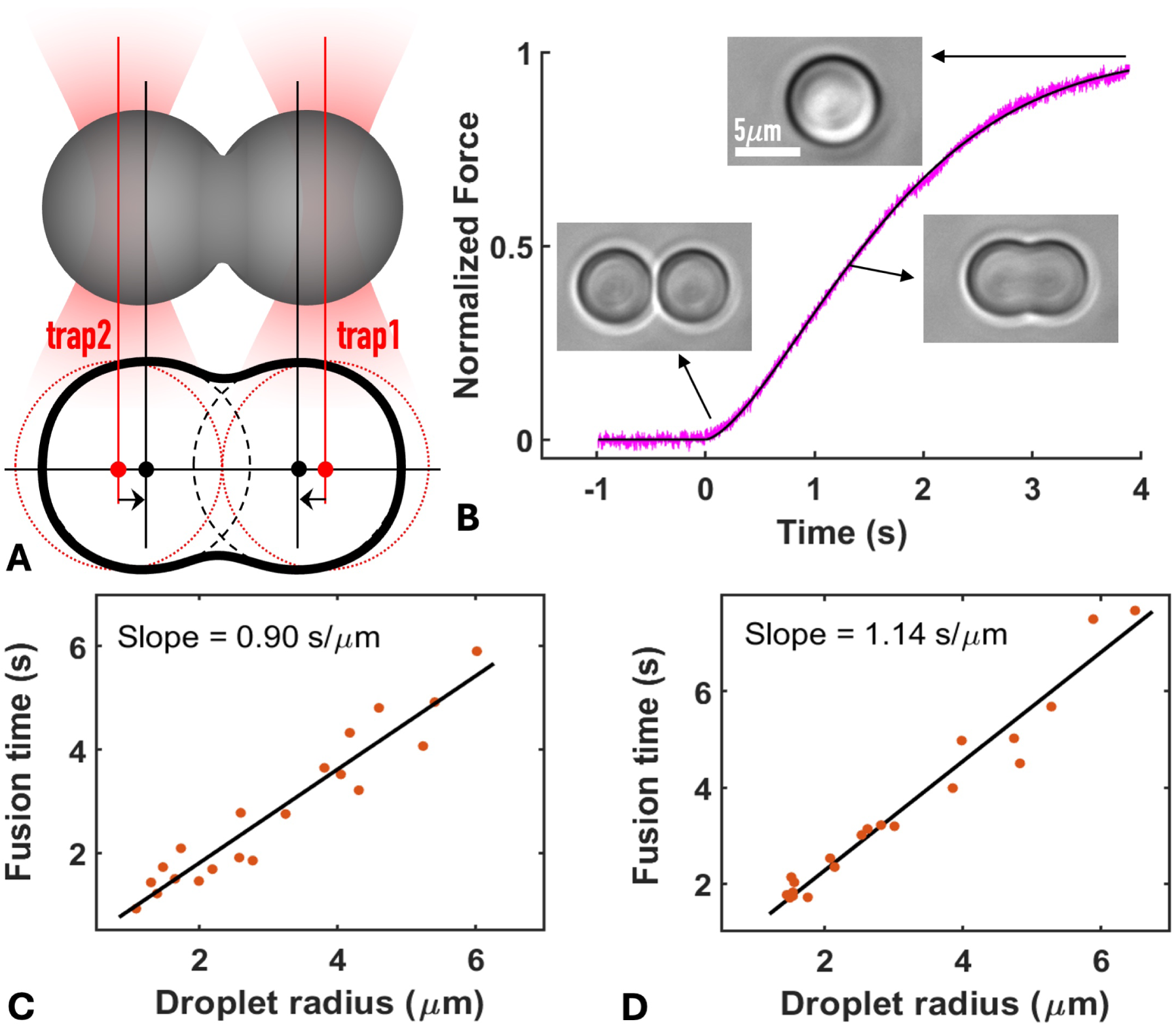
Addition of dsDNA slows down the fusion of protamine-ssDNA droplets. (A) Schematic of OT-directed droplet fusion. (B) Fusion progress curve (magenta trace) of two protamine-ssDNA droplets, monitored by the residual force reported by trap 2. Droplets were prepared by mixing 100 μM protamine with 70 μM 32-nt ssDNA. The progress curve was fit to Eq [1] (black curve) to obtain the fusion time. Brightfield images at three times are shown. (C) Plot of fusion time versus droplet radius for protamine-ssDNA droplets. (D) Corresponding plot when 1/4 of 32-nt ssDNA was replaced by 25-bp dsDNA.

## Discussion

We have demonstrated that protamine condenses a single ssDNA chain into a liquid droplet, which unravels in a sawtooth pattern that suggests the rupture of individual protamine-ssDNA bridges. The bridging forces, measured for the first time for biomolecular condensates, are 11.3 ± 4.6 pN, with unfolding lengths of 1.3 ± 0.8 µm for single bridges. Adding dsDNA raises the bridging forces to 34 ± 8 pN, reflecting stronger interactions of protamine with dsDNA than with ssDNA.

In condensing a single ssDNA chain, protamine likely brings different parts of the chain together to form nested loops (Figure 6A). Multiple protamine molecules crosslink two segments of the ssDNA chain to form a bridge. Under stretching, the protamine molecules lose contact with one ssDNA segment but stay bound to the other segment, thereby allowing ready reformation of the bridge upon retraction. When a bridge is ruptured, one half (“leading” half) of the bridge becomes loose but the other half (“lagging” half) remains inside the condensate. The resulting unfolding length is the ssDNA contour length between the leading half of this bridge and the leading half of the next bridge. In comparison, condensates that are formed on a dsDNA chain but contain a high level of ssDNA tracks unpeeled at a nick on the dsDNA are also liquid-like, with force-extension curves showing a sawtooth pattern (Figure 6B). The bridging forces are raised by the presence of dsDNA but revert to ∼10 pN by adding external ssDNA. By contrast, a high level of dsDNA leads to tangles (Figure 6C). Correspondingly, fusion is rapid for protamine-ssDNA condensates, slowed by the addition of dsDNA, and resisted when dsDNA dominates (Figure 6D).

**Figure 6.**
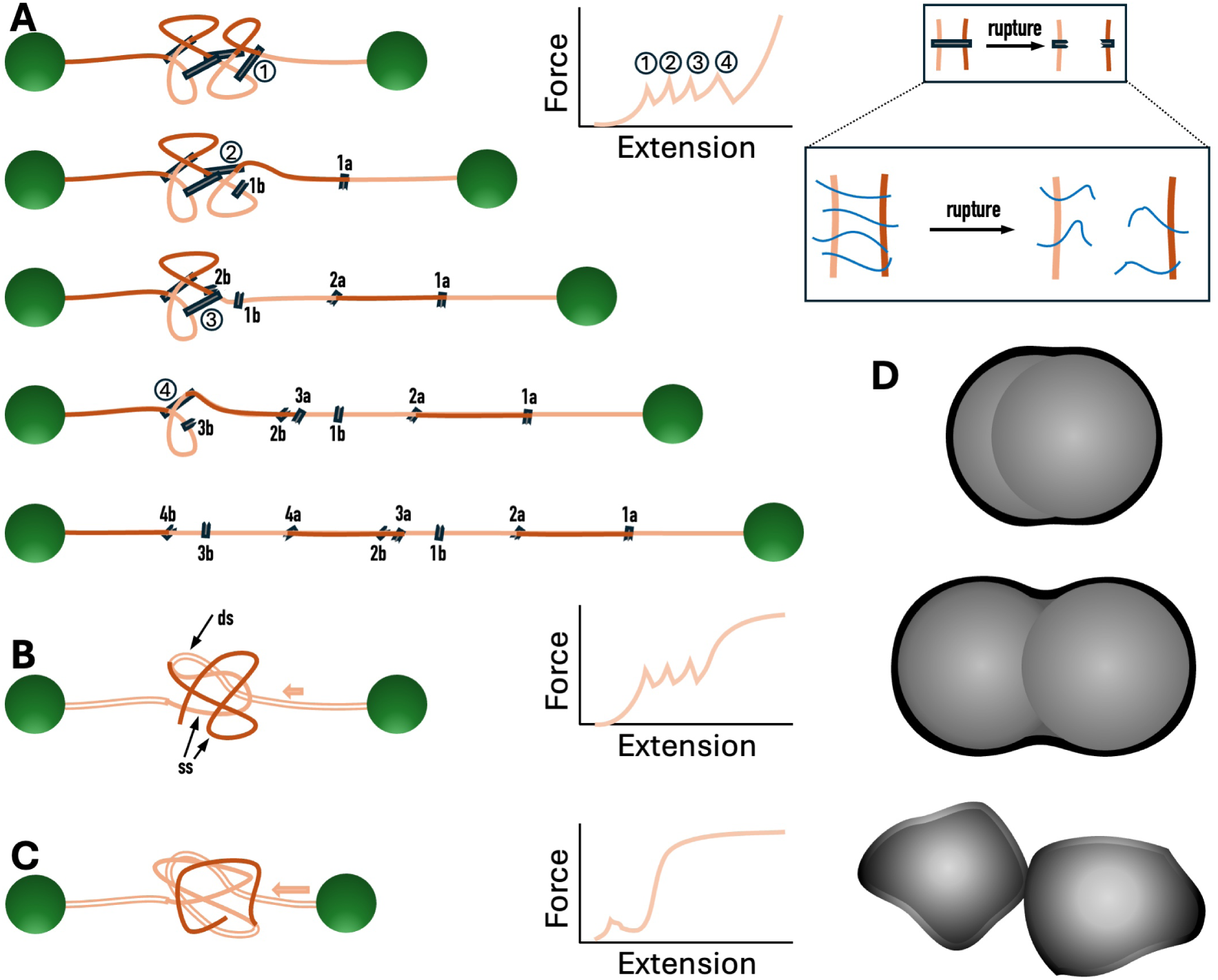
Illustration of liquid-like condensates, solid-like tangles, and intermolecular bridges. (A) Sequential unraveling of a protamine-ssDNA condensate. The leading and lagging halves of a bridge are labeled with letters “a” and “b”, respectively; the unfolded lengths of successive bridges are colored in dark and light brown. Top right: an exemplary force-extension curve and a sketch of bridge rupture. (B) A protamine-bridged condensate on a single dsDNA chain. The condensate contains ssDNA tracks unpeeled at a nick on the dsDNA as well as a relatively small amount of dsDNA drawn in by protamine (not shown). Right: the corresponding force-extension curve with a sawtooth pattern, indicating a liquid state. (C) A protamine-bridged tangle on a single dsDNA chain. The tangle contains ssDNA tracks unpeeled at a nick on the dsDNA as well as a large amount of dsDNA drawn in by protamine (not shown). Right: the corresponding force-extension curve showing an early rise to the strand-separation plateau. (D) Illustration of fusion or lack thereof for three types of condensed forms: rapid fusion for protamine-ssDNA condensates (top); slower fusion when dsDNA is added (middle); resistance of fusion when a high level of dsDNA is present (bottom).

The putatively nested loops in protamine-DNA liquid condensates are characterized by long unfolding lengths (1.3 µm), ready reformation upon rupture by force, and bridging via a disordered protein. These characteristics are distinct from observations on loops bridged by PARP1 ^27^, HMO1 ^41^, and cohesion ^42^, which are all structured proteins. In the latter cases, the unfolding lengths are short (0.1–0.3 µm) and have an exponential-like distribution, suggesting independent, randomly formed loops along the DNA molecule. Also, bridges are lost after one stretch ^41, 42^, similar to the situation with a polycation like spermine ^24^ and indicative of dissociation of bridging molecules from DNA. By contrast, due to their disordered nature, multiple protamine molecules may make up a single bridge and they may stay associated with one DNA segment after bridge rupture (Figure 6A, top right). Consequently, the sawtooth pattern is repeatedly observed over many successive stretches (Figure 2D).

The bridging forces that we have measured for protamine-DNA liquid condensates may have direct biological relevance. Protein-DNA condensates mediate transcription and regulate gene expression and DNA replication and repair ^1–12^. The forces generated by DNA-dependent motors in these processes fall in a similar range, 7.5-35 pN, as the bridging forces measured here. For example, RNA polymerases generate forces of 7.5-25 pN ^14–16^, which would be insufficient to break protamine-bridged solid tangles, leading to a plausible mechanism for global transcription silencing in the sperm nucleus. Our observation that the ssDNA-to-dsDNA ratio tunes the magnitude of bridging forces supports the idea that the bridging forces in condensates mediating DNA-related processes may be adjusted to accommodate the particular DNA-dependent motor. Uncovering other determinants for the magnitude of bridging forces will be of great interest.

When spermatids are differentiated into sperm, numerous DNA strand breaks occur to relieve supercoils as histones are removed ^22, 23^. Protamine expression also starts at this stage. Based on the fact that protamine condenses ssDNA into a liquid state (Figures 2 and 4) and that the condensates remain liquid even after a certain amount of dsDNA is added (Figure 3), we hypothesize that, in spermatids, protamine initially associates with strand breaks to form liquid condensates, which may recruit the repair machinery to restore the duplex structure. Consequently, the protamine-bridged condensates become tangles to prevent transcription in sperm.

## Materials and Methods

### Materials

Protamine sulfate (catalog # P4020) was from Sigma Aldrich. Salt was removed by dialysis ^43^ before use. SYTOX Orange and streptavidin-coated polystyrene beads (4.35 µm diameter) as a kit (catalog # 00012), biotinylated 48.5 kb λ-DNA (catalog # 00001), biotinylated 17.853-kb DNA (catalog # 00027), and biotinylated 20.452-kb DNA (catalog # 00014) were from LUMICKS. YOYO-3 (catalog # Y3606) and SYBR Gold (catalog # S11494) were from Thermo Fisher Scientific. 32-nt ssDNA, 25-bp dsDNA, and their Cy3- or Cy5-labeled versions were custom-ordered from IDT. The sequence of the 32-nt ssDNA was

5’-GAAACGCTCA GGGAAGAAGA AGATCAGCAA AG-3’

Cy3 was linked to the first nucleotide at the 5’ end. For the 25-bp dsDNA, the sequence of the first strand was

5’-ACGTTATGGC AGTCGTTAAA TTGAG-3’

Cy5 was linked to the last nucleotide at the 3’ end. Imidazole buffer (10 mM, pH 7) was prepared by dissolving imidazole (catalog # 396745000, Thermo Scientific) in Milli Q water. 150 mM KCl was added to make the working buffer. All solutions were filtered using 0.22-μm filters (catalog # SLGPR33RS, Millipore Sigma). All experiments were done at room temperature.

### Phase diagrams

Condensates were prepared by mixing imidazole buffer, KCl, ssDNA, protamine, and lastly dsDNA (if present) at desired concentrations in a microcentrifuge tube (10 µL sample volume). Aliquots of 2 μL were observed under an Olympus BX53 brightfield microscope with a 40× objective. Images were exported in tiff format and processed using ImageJ. Classification into aggregates and droplets was based on the shapes of condensates in focus. For mixtures near the boundary between aggregate and droplet regions, classification was confirmed by a brightfield camera and by OT-directed fusion on a LUMICKS C-Trap.

### Partition coefficient measurement by confocal microscopy

Partition coefficients of ssDNA and dsDNA in protamine-DNA droplets were measured using the confocal module of the C-Trap. Droplet samples were prepared by mixing either 40 µM 32-nt ssDNA or 35 µM 32-nt ssDNA and 5 µM 25bp dsDNA with 40 µM protamine in the working buffer. For measuring fluorescence intensities inside droplets, 0.01 µM of ssDNA or dsDNA was replaced with the Cy3- or Cy5-labeled version. A 10 µL aliquot of the sample was placed into a custom chamber built with a PEGylated coverslip attached to a glass slide using double-sided tape. Using an optical trap, droplets were trapped and moved around to fuse with other droplets, resulting in large droplets that were then parked on the PEGylated coverslip. After waiting for 5-10 min for any rise in temperature due to the trapping laser ^33^ to dissipate, confocal scanning of droplets was performed at 532 nm excitation for Cy3-labeled ssDNA and 638 nm excitation for Cy5-labeled dsDNA, with 0.5% laser power, 0.1 ms pixel dwell time, and 0.1 μm pixel resolution. Images were exported in tiff format and fluorescence intensities in circular regions inside droplets were read in ImageJ. To convert the fluorescence intensities into concentrations, a standard curve was determined using Cy3-labeled ssDNA or Cy5-labeled dsDNA dissolved in the working buffer. The partition coefficient was calculated by assuming that the concentration in the bulk phase was unchanged from the initial concentration (i.e., 0.01 µM). This assumption was verified by reading the fluorescence intensities in regions free of droplets when the labeled DNA concentration was increased to 0.1 µM.

### OT-directed droplet fusion

The two optical traps of the C-Trap (50:50 split of laser power) were used to trap two droplets and grow to an equal size. The overall laser power was then reduced to 3%. With the brightfield camera on, the trap-1 droplet was moved slowly to be in contact with the trap-2 droplet. Fusion then proceeded spontaneously. The force on trap 2 was recorded at a sampling rate of 78.125 kHz. The time course of the normalized trap-2 force was fit to ^39^

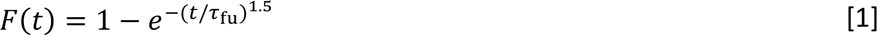

to obtain the fusion time 𝜏_fu_. The brightfield video was analyzed using ImageJ to determine the pre- and post-fusion droplet radii.

### Single-molecule force spectroscopy

Force-extension curves were acquired using the dual-trap OT module of the C-trap integrated with microfluidics (Figure S5A). The trapping laser was set to 20% overall power and a 50:50 split between trap 1 and trap 2, resulting in a stiffness of 200 pN/µm for each trap. Sample components (300-600 µL each) were loaded into syringes connected to microfluidics channels. The three main channels were filled with streptavidin-coated beads (0.004% w/v in working buffer), biotinylated DNA of different constructs in working buffer (17.853-kb DNA at 75 pg/µL, 20.452-kb DNA at 100 pg/µL, or λ-DNA at 160 pg/µL), and working buffer (10 mM Imidazole pH 7 with 150 mM KCl), respectively. The fourth, side channel was filled with protamine (10 µM in working buffer).

All channels were opened at 1 bar pressure for 2 minutes to uniformly fill the laminar flow cell with the sample components. In the laminar flow, two beads were trapped from the bead channel and then brought to the DNA channel to form a tethered configuration. After one or more tethers were formed between the two beads, the configuration was moved to the buffer channel to avoid further DNA binding. In the buffer channel, bead 2 was fixed and bead 1 was pulled until a single tether remained between the two beads. A stretching curve was acquired in the buffer channel. The tether, at either a stretched or retracted extension, was then moved to the protamine channel. After a mild flow of protamine under 0.2 bar pressure for ∼ 20 s, stretch-retract cycles were repeated until the tether broke. The pulling speed was 1 µm/s in all cases. All force-extension curves were acquired without adding dye to avoid interference unless otherwise indicated.

Further details for the three DNA constructs are as follows. For the 17.853-kb DNA, to keep it intact, a single tether was selected and stretching was limited to below 6 µm. To generate the gap version, stretching was extended to 8 µm to break weaker tethers (if present) and peel off the 5.005-knt ssDNA track between the two nicks on one strand. For either version, a stretching curve was acquired and then retracted to 2-3 µm before moving to the protamine channel. The 20.452-knt ssDNA tether was generated from the dsDNA version by overstretching. The dsDNA version had biotins at the ends of the same strand for tethering to two beads. To peel off the other strand, two-bead configurations with multiple tethers were prepared by elevating the DNA concentration, and then stretched to ∼12 µm (nearly twice the contour length of the dsDNA), at which point a single tether, with the unbiotinylated strand peeled off, remained. The ssDNA tether was retracted to ∼3 µm before moving to the protamine channel, where repeated stretch-retract cycles were performed. λ-DNA was similarly prepared at an elevated concentration to form multiple tethers; overstretching to ∼ 24 µm in buffer resulted in a single tether with unpeeled ssDNA tracks. This extended tether was then moved to the protamine channel and retracted to either 6-7 or 12-13 µm. Lastly, repeated stretch-retract cycles were performed.

### 2D confocal scans

For imaging of DNA under stretching or retraction, a dye was premixed into the protamine (10 µM) channel. The dye was SYTOX Orange (0.5 µM) for Figure 1A, SYBR Gold ( 0.5 µM) for Figure 2B, and YOYO-3 (1.2 µM) for Figures 3A and S3A. Scanning was performed with 532, 488, and 638 nm excitation for the three dyes, respectively, at 25% laser power, 0.1 ms pixel dwell time, and 0.1 μm pixel resolution. Images were processed using ImageJ.

### Force spectroscopy and 2D scans of λ-DNA mixed with external ssDNA

For this experiment (Figures S5 and S6), Cy3-labeled 32-nt ssDNA (1 μM) was loaded into the fifth, side channel (Figure S5A). The first part of the protocol was the same as that in Figure 3A, i.e., a λ-DNA tether overstretched (to ∼ 24 μm) in the buffer channel was transferred to the protamine channel (10 μM protamine and YOYO-3 1 μM) and retracted to ∼ 13 μm. After the buffer channel was opened briefly to clear out any protamine-ssDNA droplets at the entrance of the ssDNA channel, the λ-DNA tether at the fixed extension ∼ 13 μm was transferred to the ssDNA channel. After soaking with ssDNA, the λ-DNA tether at the fixed extension ∼ 13 μm was transferred back to the protamine channel. Stretch-retract cycles were finally performed until the tether broke. 2D scans were acquired simultaneously with stretch-retract cycles.

## Supporting Information

**Figure S1.**
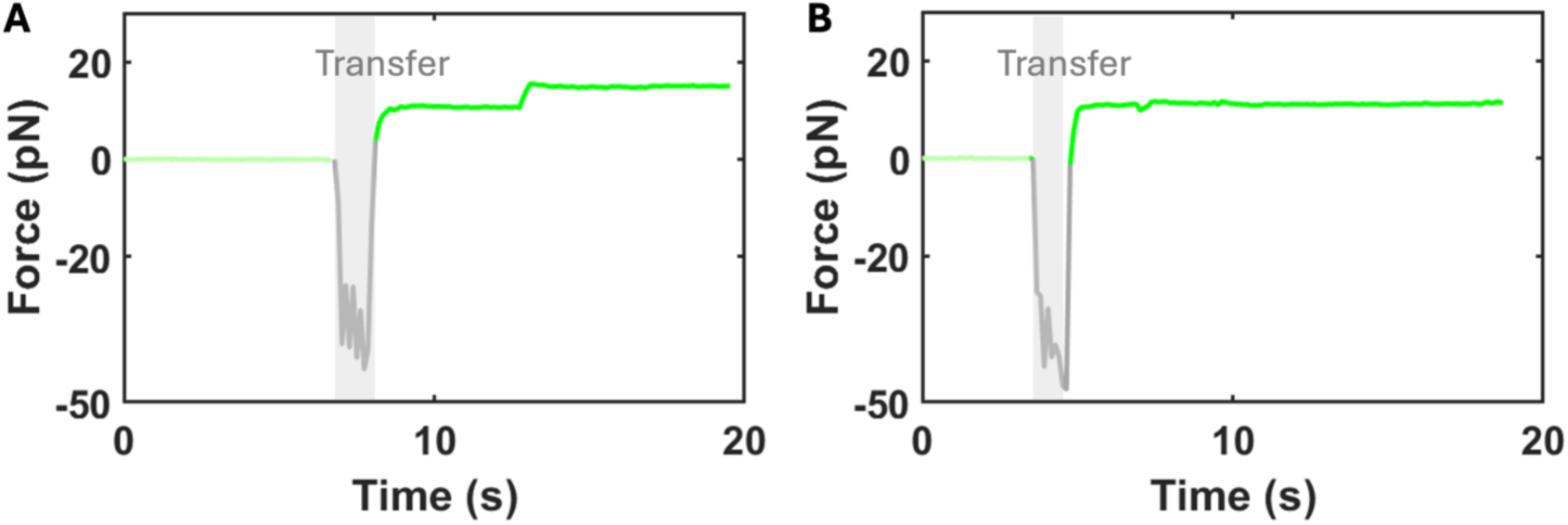
Force versus time plot illustrating a baseline shift when a retracted DNA tether is transferred from the buffer channel to the protamine channel. (A) Intact 17.853-kb DNA. (B) 20.452-knt ssDNA.

**Figure S2.**
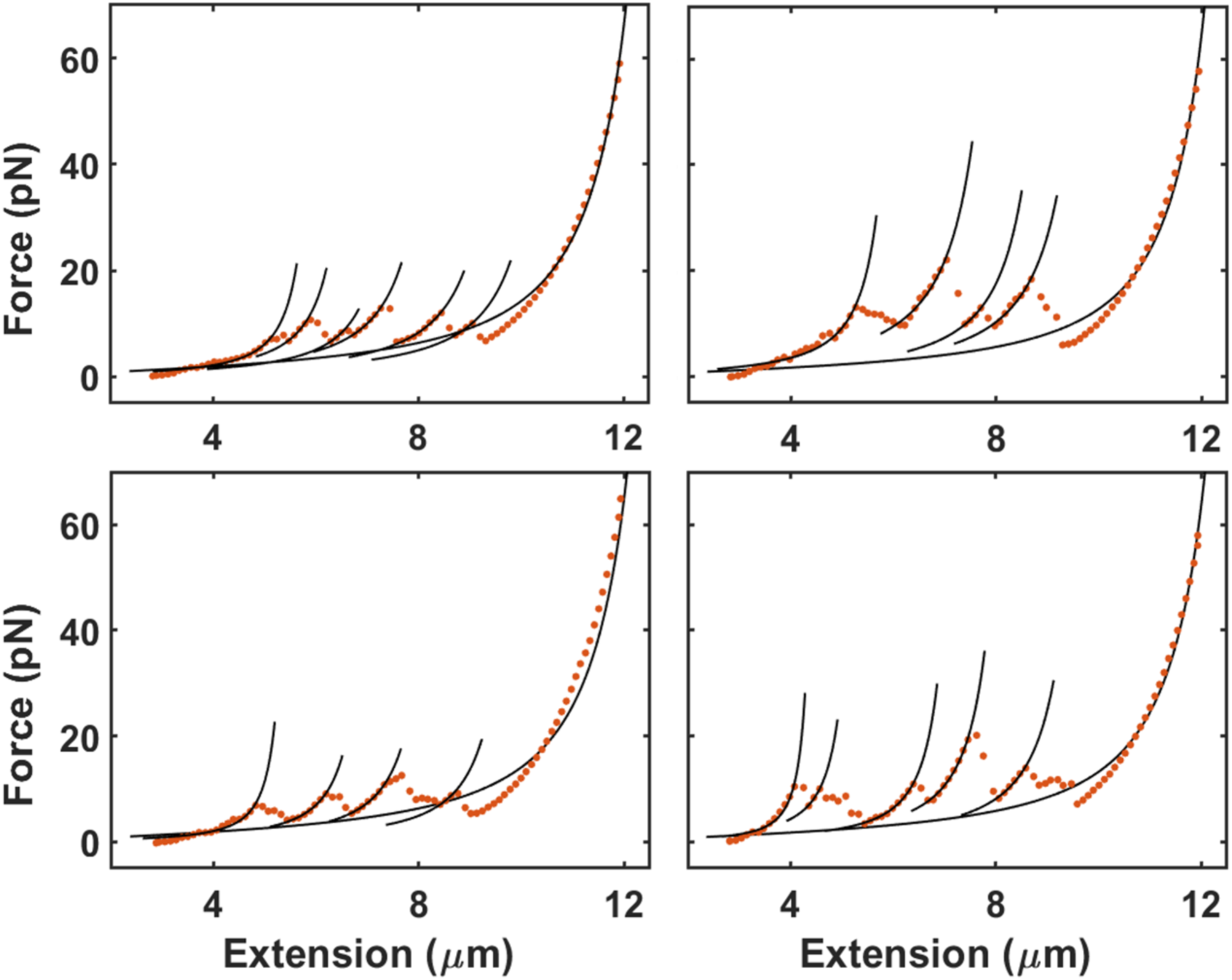
Additional force-extension curves showing unraveling of protamine-ssDNA condensates. These are similar to the one displayed in Figure 2C, but include wormlike-chain model fits to all rises in force.

**Figure S3.**
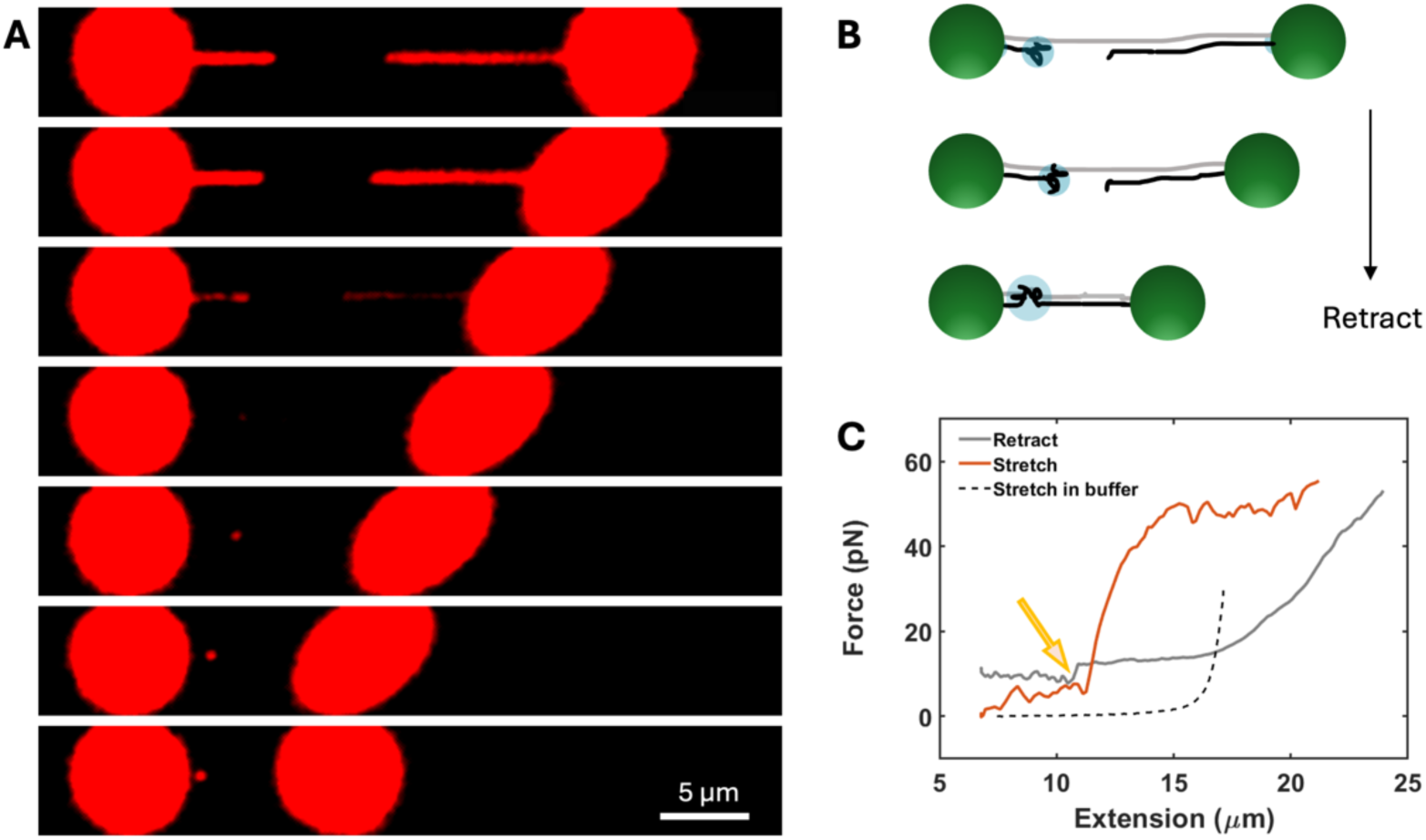
2-D scans and force-extension curves of λ-DNA containing protamine-bridged tangles initiated by overstretching. (A) Continuous 2-D scans during the retraction of overstretched λ-DNA to 7 µm. Initially, a gap representing a ssDNA track is observed; subsequently, a condensate emerges at the edge of the gap and further consolidates into a tangle. Yoyo-3 (1.2 µM) was used for staining dsDNA; the scan time per frame was 3.5 s. (B) Corresponding schematic illustrating the formation of a protamine-dsDNA-ssDNA tangle. (C) Force-extension curves for the protocol in (A) but in the absence of the dye. The retraction curve (solid gray) exhibits a sudden drop in force at ∼11 µm extension (indicated by an arrow), to a baseline that is shifted above that in the stretching curve in buffer (dashed gray). The baseline is reset to 0; the next stretch exhibits the tangling mode (orange curve).

**Figure S4.**
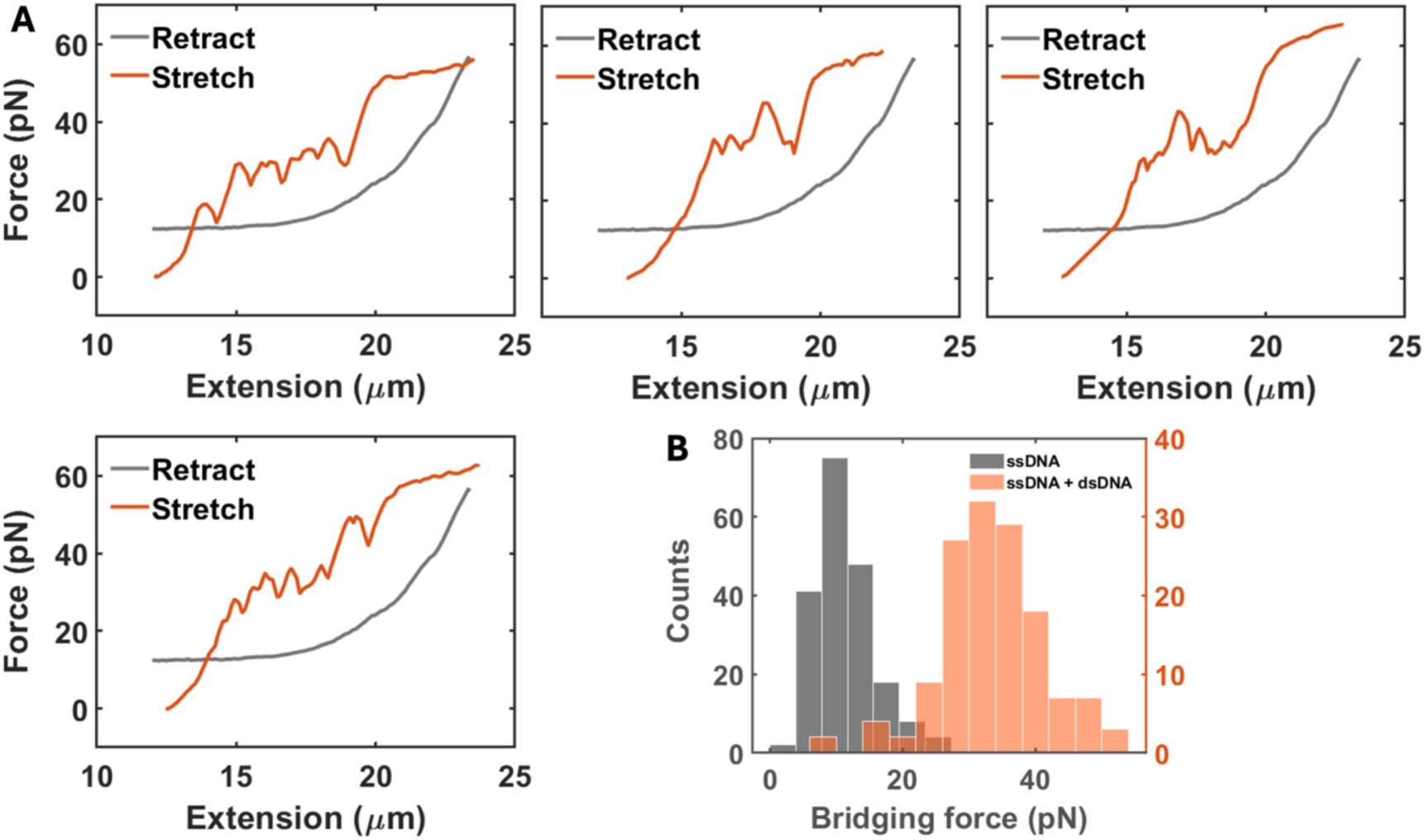
Force-extension curves of λ-DNA containing protamine-bridged condensates initiated by overstretching. (A) Stretching curves (orange) in repeated retract-stretch cycles, after the first cycle displayed in Figure 3C. The initial retraction curve (solid gray) is shown before the baseline reset. (B) Histogram of bridging forces (orange bars) from stretching curves exemplified by (A), same as that shown in Figure 3D. For comparison, the counterpart for protamine-ssDNA condensates in Figure 2E is also displayed (bray bars).

**Figure S5.**
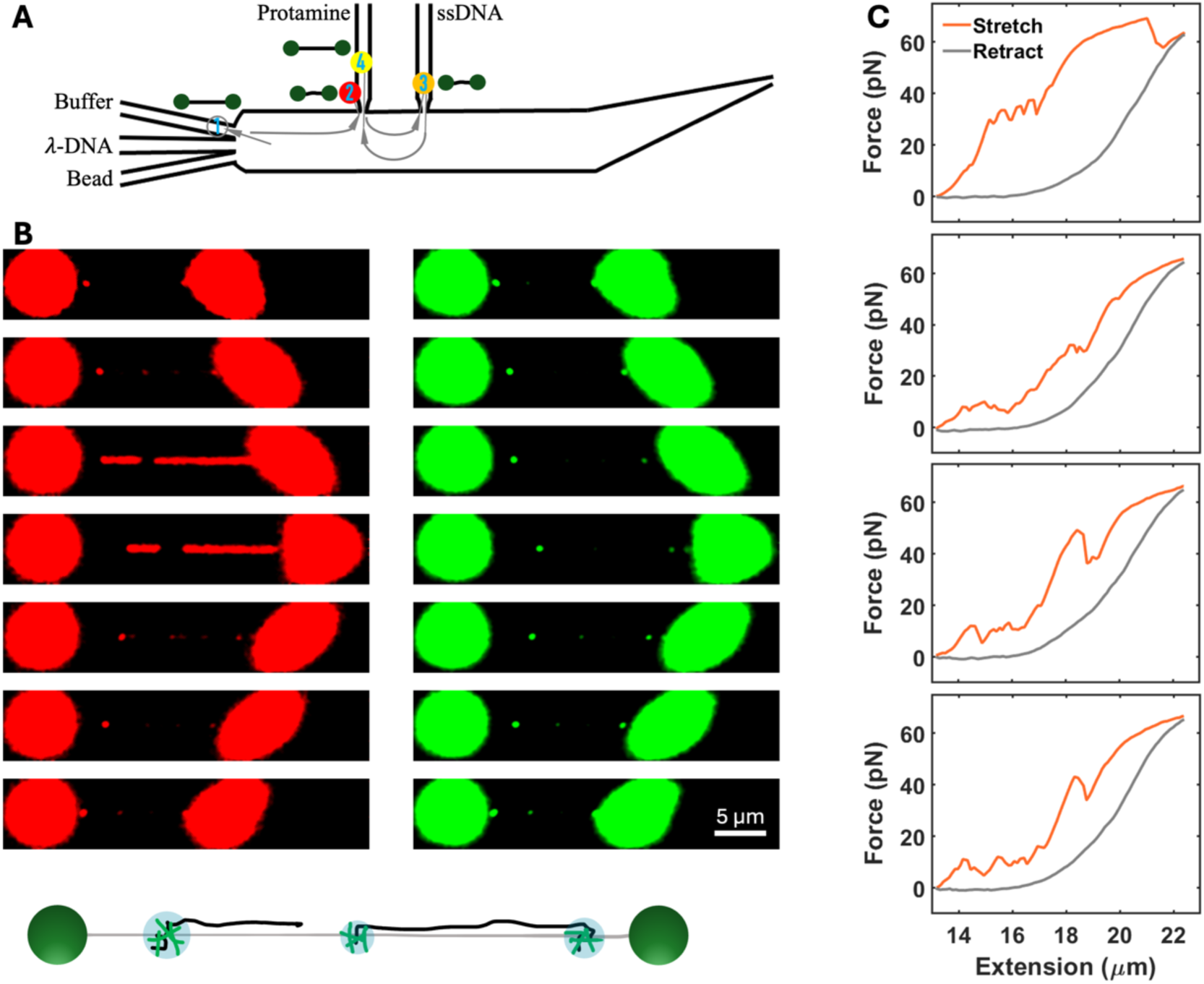
Reduction of bridging force in protamine-λ-DNA condensates by external ssDNA. (A) Illustration of the experimental protocol. (B) Continuous 2D scans in the first stretch-retract cycle of λ-DNA in the protamine channel after being soaked with 32-nt ssDNA, showing colocalization of ssDNA and dsDNA foci. A schematic is displayed at the bottom. Three prominent foci appear and persist at the edges of gaps in the λ-DNA. λ-DNA was stained by YOYO-3 (1 μM) and 32-nt ssDNA was labeled with Cy3. The scan time per frame was 3.45 s. (C) Force-extension curves in repeated stretch-retract cycles. The first stretching curve shows bridging forces at ∼35 pN and subsequent ones show a reduction of bridging forces to ∼10 pN.

**Figure S6.**
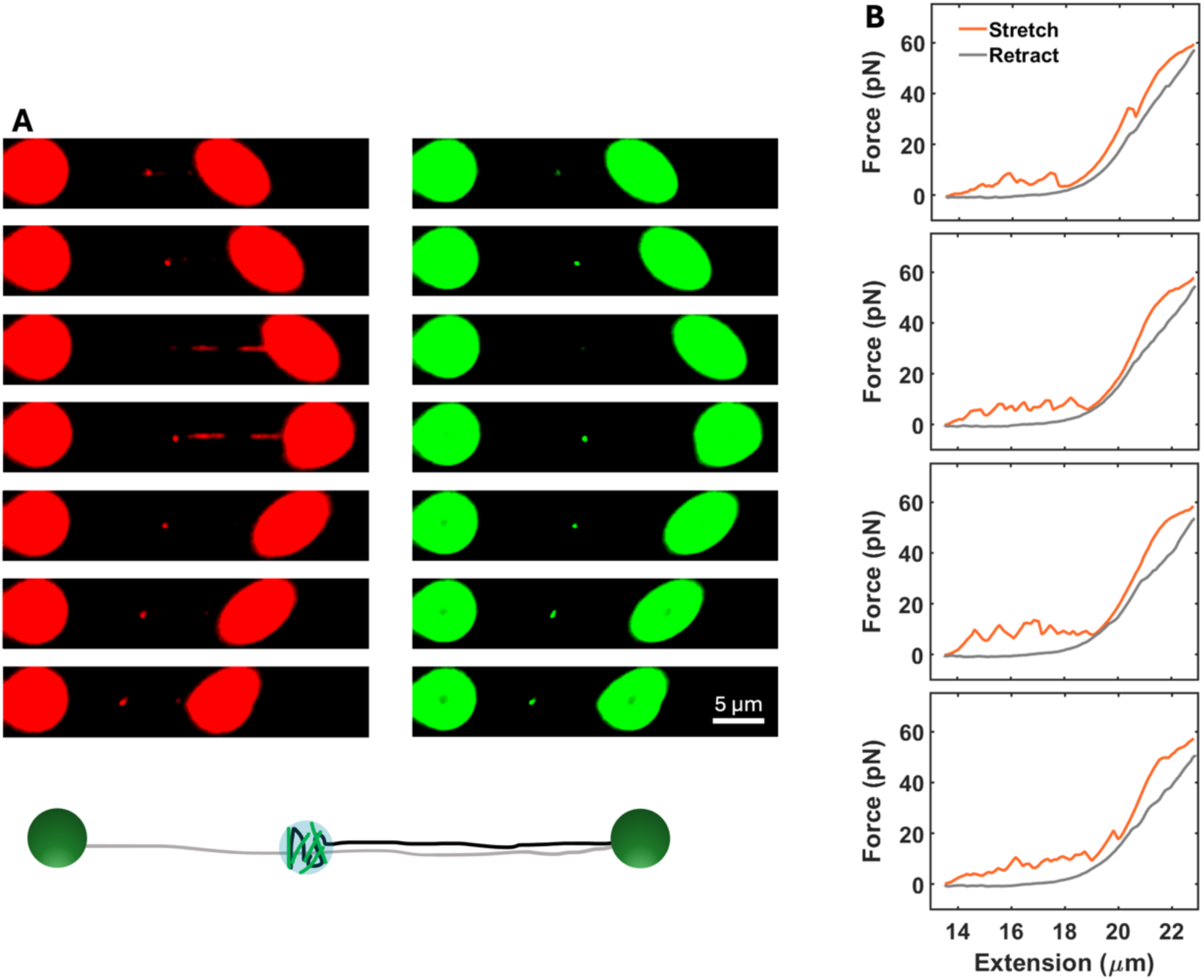
Same as Figure S5 but for a different λ-DNA tether. (A) Continuous 2D scans taken simultaneously with the first stretch-retract cycle. A schematic is displayed at the bottom. A prominent focus emerges at the edge of a major gap in the λ-DNA. A small gap also appears briefly before resealing. (B) Stretch-retract cycles. The first stretching curve already shows a reduction of bridging forces to ∼10 pN.

**Figure S7.**
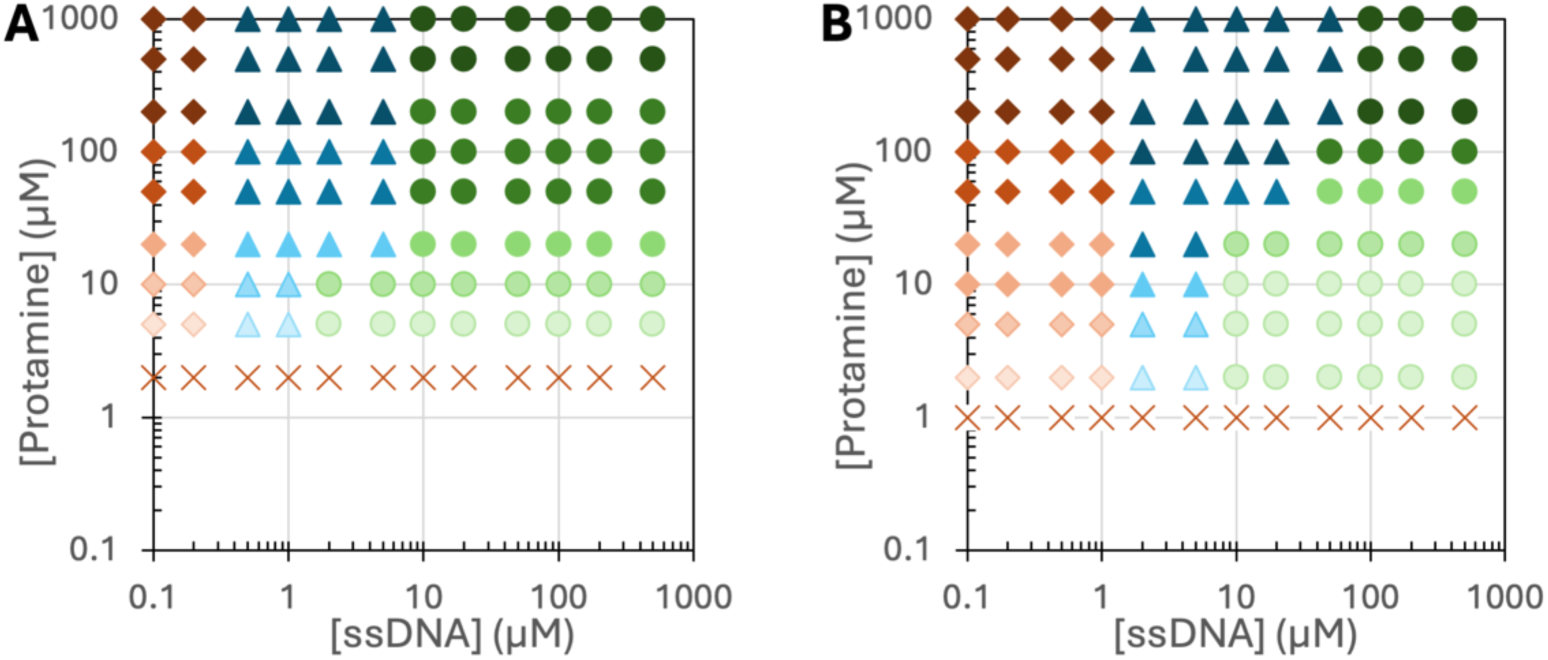
Phase diagrams of protamine-DNA mixtures. Protamine and 32-nt ssDNA were mixed in the presence of (A) 2 and (B) 10 µM 25-bp dsDNA. Cross, circles, diamonds, and triangles indicate absence of phase separation, droplets, aggregates, and mixtures of aggregates and droplets, respectively; color intensity indicates increase in condensate number and/or size.

## References

1. Cho, W. K., et al. Mediator and RNA polymerase II clusters associate in transcription-dependent condensates. Science 361, 412–415 (2018).

2. Sabari, B. R., et al. Coactivator condensation at super-enhancers links phase separation and gene control. Science 361, eaar3958 (2018).

3. Larson, A. G., et al. Liquid droplet formation by HP1α suggests a role for phase separation in heterochromatin. Nature 547, 236–240 (2017).

4. Strom, A. R., Emelyanov, A. V., Mir, M., Fyodorov, D. V., Darzacq, X., Karpen, G. H. Phase separation drives heterochromatin domain formation. Nature 547, 241–245 (2017).

5. Ji, D., et al. FOXA1 forms biomolecular condensates that unpack condensed chromatin to function as a pioneer factor. Mol Cell 84, 244–260.e247 (2024).

6. Parker, M. W., et al. A new class of disordered elements controls DNA replication through initiator self-assembly. eLife 8, e48562 (2019).

7. Kilic, S., et al. Phase separation of 53BP1 determines liquid-like behavior of DNA repair compartments. EMBO J 38, e101379 (2019).

8. Singatulina, A. S., et al. PARP-1 Activation Directs FUS to DNA Damage Sites to Form PARG-Reversible Compartments Enriched in Damaged DNA. Cell Rep 27, 1809–1821 e1805 (2019).

9. Oshidari, R., et al. DNA repair by Rad52 liquid droplets. Nat Commun 11, 695 (2020).

10. Claeys Bouuaert, C., et al. DNA-driven condensation assembles the meiotic DNA break machinery. Nature 592, 144–149 (2021).

11. Spegg, V., et al. Phase separation properties of RPA combine high-affinity ssDNA binding with dynamic condensate functions at telomeres. Nat Struct Mol Biol 30, 451–462 (2023).

12. Chappidi, N., et al. PARP1-DNA co-condensation drives DNA repair site assembly to prevent disjunction of broken DNA ends. Cell 187, 945–961 e918 (2024).

13. Bustamante, C., Bryant, Z., Smith, S. B. Ten years of tension: single-molecule DNA mechanics. Nature 421, 423–427 (2003).

14. Wang, M. D., Schnitzer, M. J., Yin, H., Landick, R., Gelles, J., Block, S. M. Force and Velocity Measured for Single Molecules of RNA Polymerase. Science 282, 902–907 (1998).

15. John, R., Wuite, G. J. L., Landick, R., Bustamante, C. Single-Molecule Study of Transcriptional Pausing and Arrest by *E. coli* RNA Polymerase. Science 287, 2497–2500 (2000).

16. Galburt, E. A., et al. Backtracking determines the force sensitivity of RNAP II in a factor-dependent manner. Nature 446, 820–823 (2007).

17. Maier, B., Bensimon, D., Croquette, V. Replication by a single DNA polymerase of a stretched single-stranded DNA. Proc Natl Acad Sci U S A 97, 12002–12007 (2000).

18. Wuite, G. J. L., Smith, S. B., Young, M., Keller, D., Bustamante, C. Single-molecule studies of the effect of template tension on T7 DNA polymerase activity. Nature 404, 103–106 (2000).

19. Manosas, M., Perumal, S. K., Bianco, P. R., Ritort, F., Benkovic, S. J., Croquette, V. RecG and UvsW catalyse robust DNA rewinding critical for stalled DNA replication fork rescue. Nature Communications 4, 2368 (2013).

20. Rathke, C., Baarends, W. M., Awe, S., Renkawitz-Pohl, R. Chromatin dynamics during spermiogenesis. Biochim Biophys Acta 1839, 155–168 (2014).

21. Kitaoka, M., Yamashita, Y. M. Running the gauntlet: challenges to genome integrity in spermiogenesis. Nucleus 15, 2339220 (2024).

22. Marcon, L., Boissonneault, G. Transient DNA strand breaks during mouse and human spermiogenesis new insights in stage specificity and link to chromatin remodeling. Biol Reprod 70, 910–918 (2004).

23. Rathke, C., Baarends, W. M., Jayaramaiah-Raja, S., Bartkuhn, M., Renkawitz, R., Renkawitz-Pohl, R. Transition from a nucleosome-based to a protamine-based chromatin configuration during spermiogenesis in Drosophila. J Cell Sci 120, 1689–1700 (2007).

24. Ahlawat, V., Dhiman, A., Mudiyanselage, H. E., Zhou, H. X. Protamine-Mediated Tangles Produce Extreme Deoxyribonucleic Acid Compaction. J Am Chem Soc 146, 30668–30677 (2024).

25. Quail, T., et al. Force generation by protein–DNA co-condensation. Nat Phys 17, 1007–1012 (2021).

26. Morin, J. A., et al. Sequence-dependent surface condensation of a pioneer transcription factor on DNA. Nat Phys 18, 271–276 (2022).

27. Bell, N. A. W., et al. Single-molecule measurements reveal that PARP1 condenses DNA by loop stabilization. Sci Adv 7, eabf3641 (2021).

28. Keenen, M. M., et al. HP1 proteins compact DNA into mechanically and positionally stable phase separated domains. eLife 10, e64563 (2021).

29. Nguyen, T., et al. Chromatin sequesters pioneer transcription factor Sox2 from exerting force on DNA. Nat Commun 13, 3988 (2022).

30. Gien, H., et al. HIV-1 Nucleocapsid Protein Binds Double-Stranded DNA in Multiple Modes to Regulate Compaction and Capsid Uncoating. Viruses 14, 235 (2022).

31. Leicher, R., et al. Single-stranded nucleic acid binding and coacervation by linker histone H1. Nat Struct Mol Biol 29, 463–471 (2022).

32. Renger, R., et al. Co-condensation of proteins with single- and double-stranded DNA. Proc Natl Acad Sci U S A 119, e2107871119 (2022).

33. Pokhrel, P., Jonchhe, S., Pan, W., Mao, H. Single-Molecular Dissection of Liquid– Liquid Phase Transitions. J Am Chem Soc 145, 17143–17150 (2023).

34. Vieregg, J. R., et al. Oligonucleotide–Peptide Complexes: Phase Control by Hybridization. J Am Chem Soc 140, 1632–1638 (2018).

35. Shakya, A., King, J. T. DNA Local-Flexibility-Dependent Assembly of Phase-Separated Liquid Droplets. Biophys J 115, 1840–1847 (2018).

36. Marko, J. F., Siggia, E. D. Stretching DNA. Macromolecules 28, 8759–8770 (1995).

37. King, G. A., Gross, P., Bockelmann, U., Modesti, M., Wuite, G. J. L., Peterman, E. J. G. Revealing the competition between peeled ssDNA, melting bubbles, and S-DNA during DNA overstretching using fluorescence microscopy. Proc Natl Acad Sci U S A 110, 3859–3864 (2013).

38. Zhang, Y., Prasad, R., Su, S., Lee, D., Zhou, H. X. Amino acid-dependent phase equilibrium and material properties of tetrapeptide condensates. Cell Rep Phys Sci 5, (2024).

39. Ghosh, A., Zhou, H. X. Determinants for Fusion Speed of Biomolecular Droplets. Angew Chem Int Ed Engl 59, 20837–20840 (2020).

40. Ghosh, A., Kota, D., Zhou, H. X. Shear relaxation governs fusion dynamics of biomolecular condensates. Nat Commun 12, 5995 (2021).

41. Murugesapillai, D., et al. DNA bridging and looping by HMO1 provides a mechanism for stabilizing nucleosome-free chromatin. Nucleic Acids Res 42, 8996–9004 (2014).

42. Gutierrez-Escribano, P., et al. A conserved ATP- and Scc2/4-dependent activity for cohesin in tethering DNA molecules. Sci Adv 5, eaay6804 (2019).

43. Kota, D., Prasad, R., Zhou, H. X. Adenosine Triphosphate Mediates Phase Separation of Disordered Basic Proteins by Bridging Intermolecular Interaction Networks. J Am Chem Soc 146, 1326–1336 (2024).

